# Asexual Adaptation Drives Transient and Reversible Changes in Mating Efficiency

**DOI:** 10.64898/2026.05.14.725142

**Authors:** Prachitha Nagendra, Deesha Kansara, Aditi Sriram, Supreet Saini

## Abstract

The evolution of reproductive isolation is central to speciation, yet the earliest stages of this process remain poorly understood. In particular, it is unclear how rapidly barriers to mating arise during adaptation, whether they accumulate predictably, and how they depend on ecological context. Here, we investigate the evolution of mating efficiency during prolonged asexual adaptation in diploid *Saccharomyces cerevisiae*. Twelve replicate populations were evolved for 1200 generations in two distinct carbon environments, glucose and galactose, under strictly asexual conditions. At regular intervals, we induced sporulation and quantified mating efficiency using three complementary assays: within-population crosses, crosses between populations evolved in different environments, and crosses between evolved populations and the ancestral strain. We find that mating efficiency evolves during asexual adaptation, with outcomes that depend strongly on the environment. While glucose-evolved populations remain largely stable, galactose-evolved populations exhibit a reversible decline. Overall, changes in mating efficiency are dynamic, heterogeneous, and often transient, with evidence for both intrinsic reductions in mating competence and context-dependent incompatibilities between populations. Together, these results show that asexual adaptation can generate rapid but non-monotonic changes in mating compatibility. Early reductions in mating efficiency are heterogeneous, environment-dependent, and often reversible, and do not accumulate into stable reproductive isolation over the timescale examined. Our findings suggest that the initial stages of divergence are characterized by dynamic and contingent perturbations of reproductive traits, rather than a steady progression toward speciation.

## Introduction

The evolution of reproductive isolation is a central problem in evolutionary biology, yet the earliest stages of this process remain poorly understood (1-5). In particular, it is unclear how quickly barriers to mating can arise, whether they accumulate predictably over time, and to what extent they depend on ecological context (1, 6-10). While classical models of speciation emphasize the gradual buildup of incompatibilities between diverging populations, increasing evidence suggests that early changes in reproductive traits may be more dynamic, contingent, and reversible than traditionally assumed (4,10-14).

Experimental evolution provides a powerful framework to study the emergence of reproductive isolation under controlled conditions (8, 12, 15-17). Microbial systems, in particular, enable the tracking of evolutionary trajectories across hundreds to thousands of generations, while maintaining replicate populations evolving under identical or contrasting environments (16, 18, 19). However, most experimental evolution studies focus on fitness and adaptation, with comparatively less attention given to the evolution of reproductive traits (8, 12, 16, 18-20). As a result, the extent to which adaptation in different environments influences mating compatibility and whether such effects are consistent across replicate populations remains an open question (10, 21, 22).

In facultatively sexual organisms such as budding yeast, mating is a multistep process involving pheromone signaling, cell cycle arrest, and cell fusion (23-26). These processes are governed by regulatory networks that are not directly under selection during prolonged asexual growth (27-31). Consequently, mutations that accumulate during asexual evolution may have pleiotropic effects on mating efficiency, defined here as the proportion of cells that successfully form diploids upon mixing compatible haploid partners (32-36). Such effects could lead to reduced mating performance within populations or to incompatibilities between populations that have adapted to different environments (4, 5, 8, 12, 37). At the same time, compensatory evolution or network-level robustness may buffer or reverse these changes, resulting in non-monotonic trajectories of reproductive traits (38-42).

Several key questions therefore arise. First, how does mating efficiency evolve during extended periods of asexual adaptation? (8, 12) Second, does adaptation to distinct environments generate reduced compatibility between populations, consistent with the emergence of pre-zygotic barriers? (21) Third, are such changes predictable and parallel across replicate populations, or do they depend on lineage-specific evolutionary trajectories? (13, 19) Finally, do reductions in mating efficiency reflect intrinsic changes in mating competence, or do they primarily arise from interactions between independently evolved genomes? (10)

Here, we address these questions using experimental evolution in diploid *Saccharomyces cerevisiae*. We evolved replicate populations for 1200 generations in two distinct carbon environments, glucose and galactose, under strictly asexual conditions. Adaptation in these environments has been well studied and has been a focus of large number of experimental studies (43-47). At regular intervals,we induced sporulation and quantified mating efficiency in three complementary assays: (i) within-population crosses between haploids derived from the same evolved line, (ii) crosses between populations evolved in different environments, and (iii) crosses between evolved populations and the ancestral strain. This design allows us to disentangle intrinsic changes in mating competence from pairwise incompatibilities and to assess the repeatability of evolutionary outcomes across replicate populations.

By tracking mating efficiency across environments, time, and defined mating contexts, our study provides a detailed view of how reproductive traits evolve during asexual adaptation. In doing so, we aim to clarify whether early changes in mating compatibility follow a directional trajectory toward reproductive isolation or instead reflect dynamic and heterogeneous evolutionary processes.

## Results

### Mating efficiency evolves during asexual adaptation in an environment-dependent manner

To assess how asexual evolution influences mating performance, we sporulated diploid populations evolved in glucose or galactose at 300-generation intervals and quantified mating efficiency between haploids derived from the same population (**Figures 1**).

**Figure 1.**
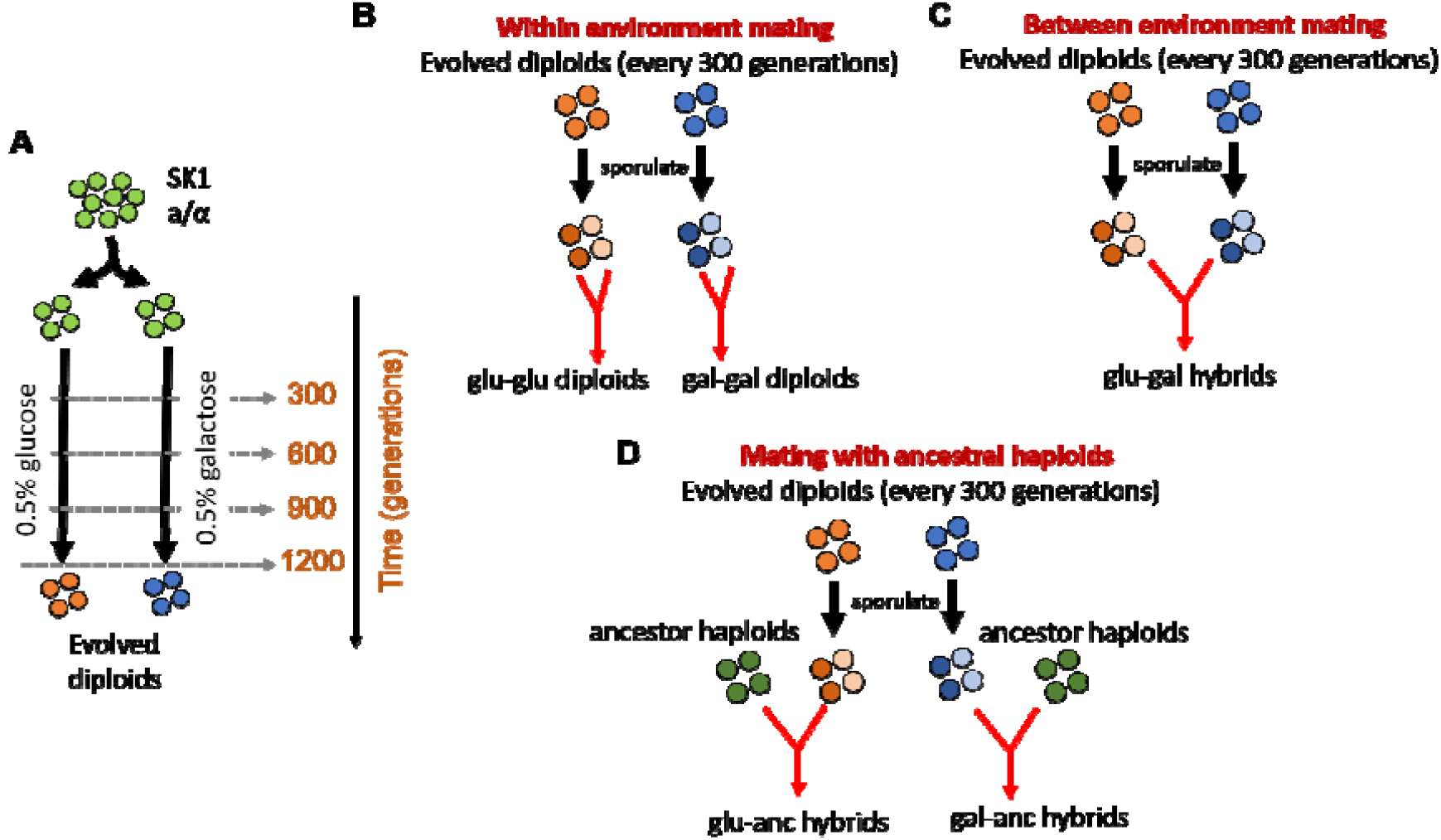
Experimental design. **(A)** Isogenic population of *S. cerevisiae* was evolved asexually in glucose and galactose until 1200 generations (48). Mating efficiency assays were performed every 300 generations. At each time point, evolved populations were sporulated to obtain haploids, which were then used in the following mating assays: **(B)** Within environment mating. Haploids derived from a population at different time points were mated, **(C)** Between environment mating. Haploids obtained from populations evolved in the two environments at different time points were crossed, and **(D)** Mating with ancestor haploids. Haploids obtained from evolved diploids were crossed with the ancestral haploids.

In glucose-evolved populations, through the course of the experiment, mating efficiency did not change statistically, as compared to the ancestor. Across all time-points (300–900 generations), median mating efficiency values were maintained at approximately 80–85%, with only modest fluctuations (**Figure 2A**). However, at 1200 generations, we note a statistically significant decrease in the mating efficiency of the evolved haploids **(Supplement Figure S1)**. Although some individual measurements fell below this range, there was no systematic decline over time, indicating that adaptation in glucose does not substantially impair mating performance on average. This was somewhat surprising since effective mating was not selected for the period of adaptation. Despite there being no statistical change in the mating efficiency with time, we did observe an increase in the variance of mating efficiency with time. As shown in **Figure 2B**, the range of mating efficiency values observed during adaptation in glucose increased with time.

**Figure 2.**
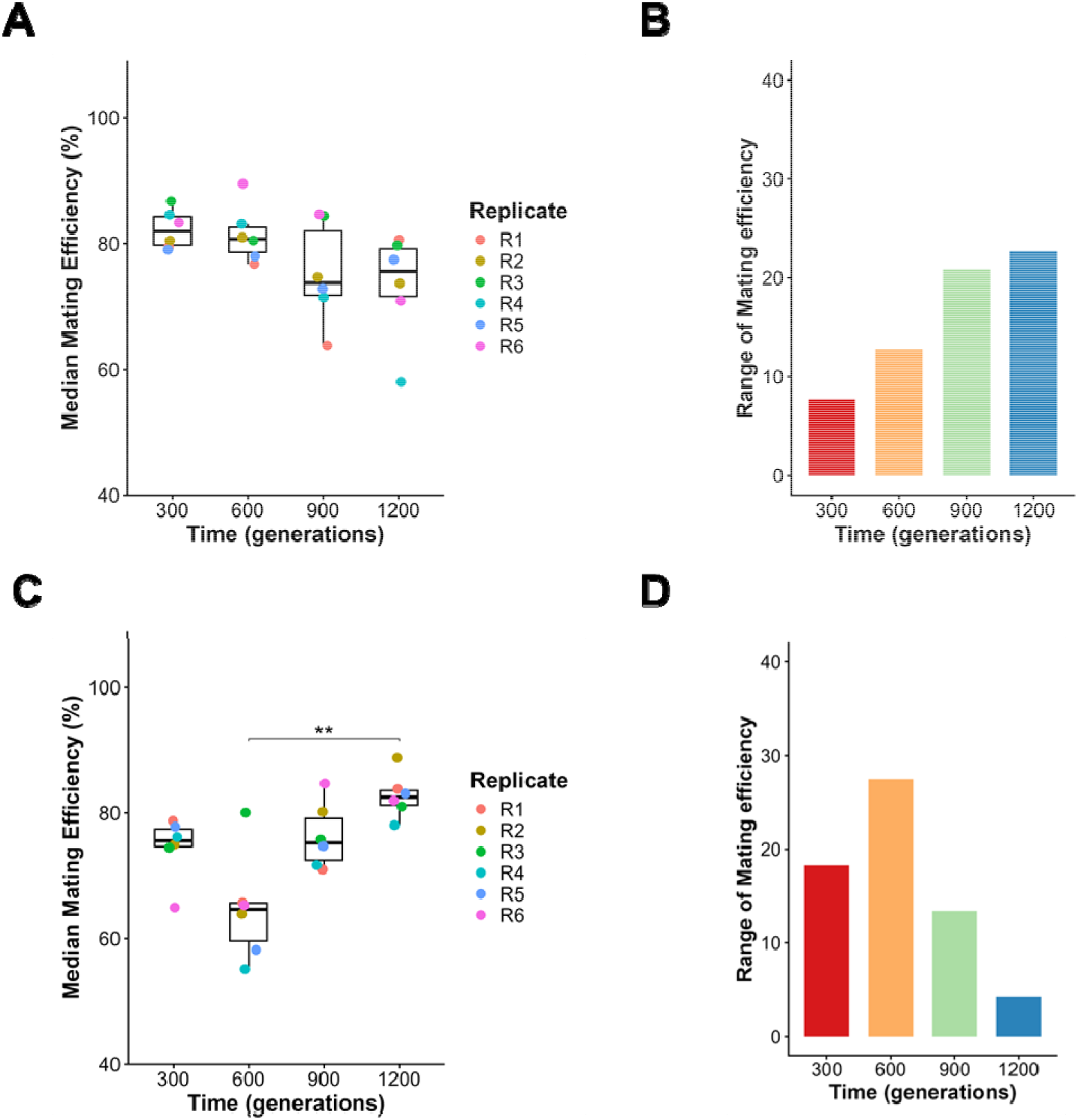
Mating efficiency of the evolved haploids changes with time. **(A)** The glucose-evolved population was sporulated and haploids from the same population were allowed to mate with one another to assess the effect of asexual evolution on mating efficiency. Median mating efficiency was calculated for every replicate at each time point. Across all four time points, median mating efficiency remained consistently high, ranging from 70-85%, with no significant differences in the mean mating efficiency among time (Kruskal Wallis Test, p > 0.05). **(B)** Range of the six median mating efficiency values increases with time of evolution in glucose. **(C)** The galactose-evolved population was sporulated and haploids from the same population were allowed to mate with one another to assess the effect of asexual evolution on mating efficiency. Median mating efficiency was calculated for every replicate at each time point. Unlike glucose-evolved populations, galactose-evolved populations exhibited a temporal effect on mating efficiency (Kruskal Wallis test, p<0.01). At the early time point, the mating efficiency was high and comparable to that of glucose-evolved populations. However, at 600 generations, mating efficiency showed a decline, ranging from 55-65%, followed by gradually recovery by 1200 generations. Pairwise comparisons using Dunn’s test indicated significant differences between 600 and 1200 generations (p<0.01). **(D)** Range of the six median mating efficiency values changes non-monotonically with time of evolution in glucose.

In contrast, galactose-evolved populations exhibited a pronounced and temporal reduction in mating efficiency (**Figure 2C**). At 300 generations, mating efficiency was comparable to that of glucose-evolved populations (∼80%). This mating efficiency was, however, lower than the ancestor mating efficiency **(Supplement Figure S1)**. However, by 600 generations, median mating efficiency declined markedly to ∼60–65%. Over the next 600 generations, i.e., by 1200 generations, mating efficiency recovered to values statistically identical to the ancestor’s mating efficiency. Interestingly, as the mating efficiency of the galactose-evolved lines decreased, the range of median mating efficiency between the six lines increased. As shown in **Figure 2D**, the range of mating efficiency was maximum at 600 generations (when the mating efficiency was the minimum). In addition, at 1200 generations, the range of mating efficiency was minimum when the mating efficiency was maximum (although not statistically significant).

These results demonstrate that mating efficiency evolves during asexual adaptation, with both the magnitude and temporal dynamics of change dependent on the selective environment. In particular, adaptation to galactose is associated with a substantial but reversible reduction in mating performance. From the context of studying emergence of reproductive barriers, asexual adaptation in an environment for a duration of ∼1200 generations does not significantly impair a population’s ability to mate.

### Cross-environment mating reveals transient reductions in hybrid mating efficiency

To test whether adaptation to different environments leads to reduced mating efficiency between populations, we performed crosses between haploids derived from glucose- and galactose-evolved lines at corresponding time points (**Figure 3A**).

**Figure 3.**
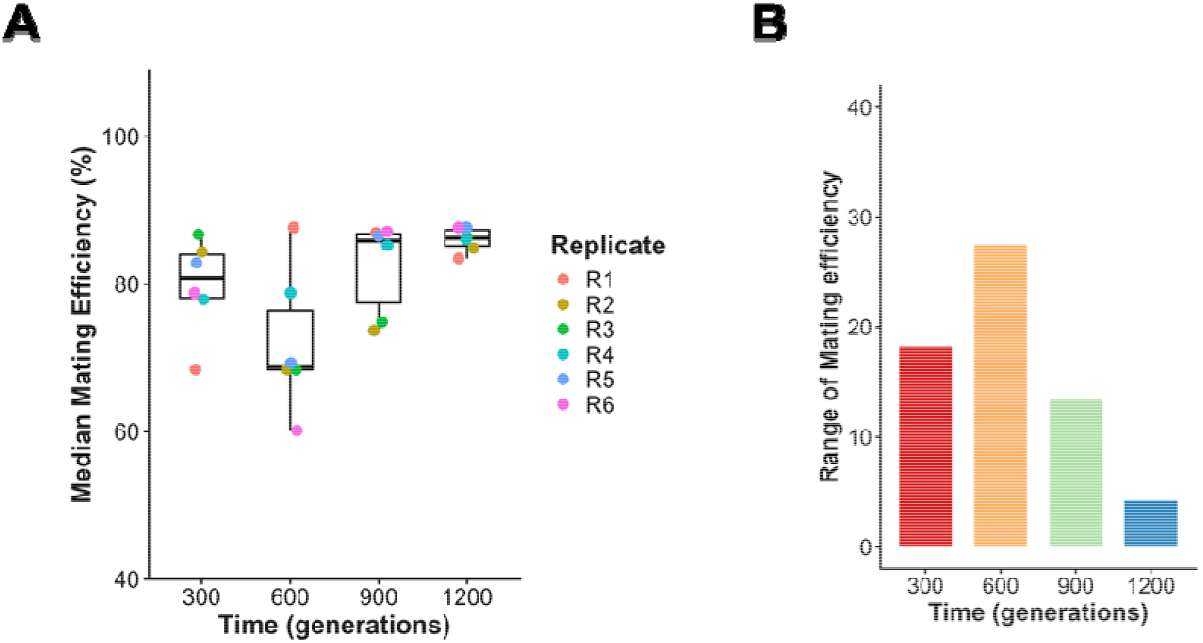
Mating efficiency between haploids derived from glucose- and galactose-evolved populations. **(A)** Glucose- and galactose-evolved population were sporulated and haploids from the two environments were crossed to study the effect of asexual evolution on reproductive isolation. A transient reduction in mating efficiency was observed in 1200 generations, however it was not statistically significant. Hybrid mating efficiency remained high, ranging from 70-85%, at both early and later time points. A significant reduction was observed only at 600 generations, although substantial variability was present among replicates. **(B)** Range of the six median mating efficiency in glu-gal hybrids is maximum at 600 generations and is the least at 1200 generations.

Hybrid mating efficiency was relatively high at early time-points (300 generations), with values comparable to those exhibited by the ancestor and those observed within environments (Figure 2). However, at 600 generations, a reduction in mating efficiency was observed across hybrid crosses. Although this reduction was statistically insignificant, it was ear-marked with a large increase in the range of the six median values for the six independent lines.

Importantly, the decrease in hybrid mating efficiency was not sustained. At later time-points (900 and 1200 generations), mating efficiency increased again, approaching levels similar to those observed at earlier stages of the experiment. The recovery of the mating efficiency was accompanied by a reduction in variability between the six lines. This result is shown in **Figure 3B** where the range of median values of the mating efficiency is maximum at 600 generations, and then reduces by 1200 generations. This trend mirrors the dynamics of range of mating efficiency values exhibited by galactose-evolved lines (Figure 2D). As seen in Figure 2D, in the case of hybrid matings, the range of the median values of the mating efficiency is inversely correlated with the mean of the median values of the mating efficiency.

On comparing the mating efficiency within a line (Figure 2) vs. the mating efficiency between lines (Figure 3), we note that the hybrid mating efficiency is greater than the within-line mating efficiency in all cases except one (**Supplement Figure S2**). The only exception to the above is at 600 generations, when the mating efficiency of glu-glu lines is greater than that between the glu-gal lines. These results show that evolution of pre-zygotic barrier is a temporal and ephemeral phenomenon. For its effects to lead to biological consequences, it likely needs to be accompanied with other forces/phenomenon. Transient emergence of pre-zygotic barriers, by themselves, are unlikely to persist.

### Crosses to the ancestral strain reveal intrinsic changes in mating competence

To determine whether reductions in mating efficiency reflect pairwise incompatibilities between evolved populations or intrinsic changes in mating ability, we crossed evolved haploids to haploids derived from the ancestral strain.

In glucose-evolved populations (**Figure 4A**), mating efficiency with the ancestor reduced slightly (statistically insignificant) in the first 600 generations. However, in the subsequent 600 generations,the mating efficiency between the evolved lines and the ancestor haploids increased to ancestor levels of mating efficiency. Mating efficiency of glucose-evolved with ancestor, however,is always significantly lower than that of ancestral mating efficiency **(Supplement Figure S3)**. This was accompanied by relatively high variance of mating efficiency between the lines at the four time points (**Figure 4B**).

**Figure 4.**
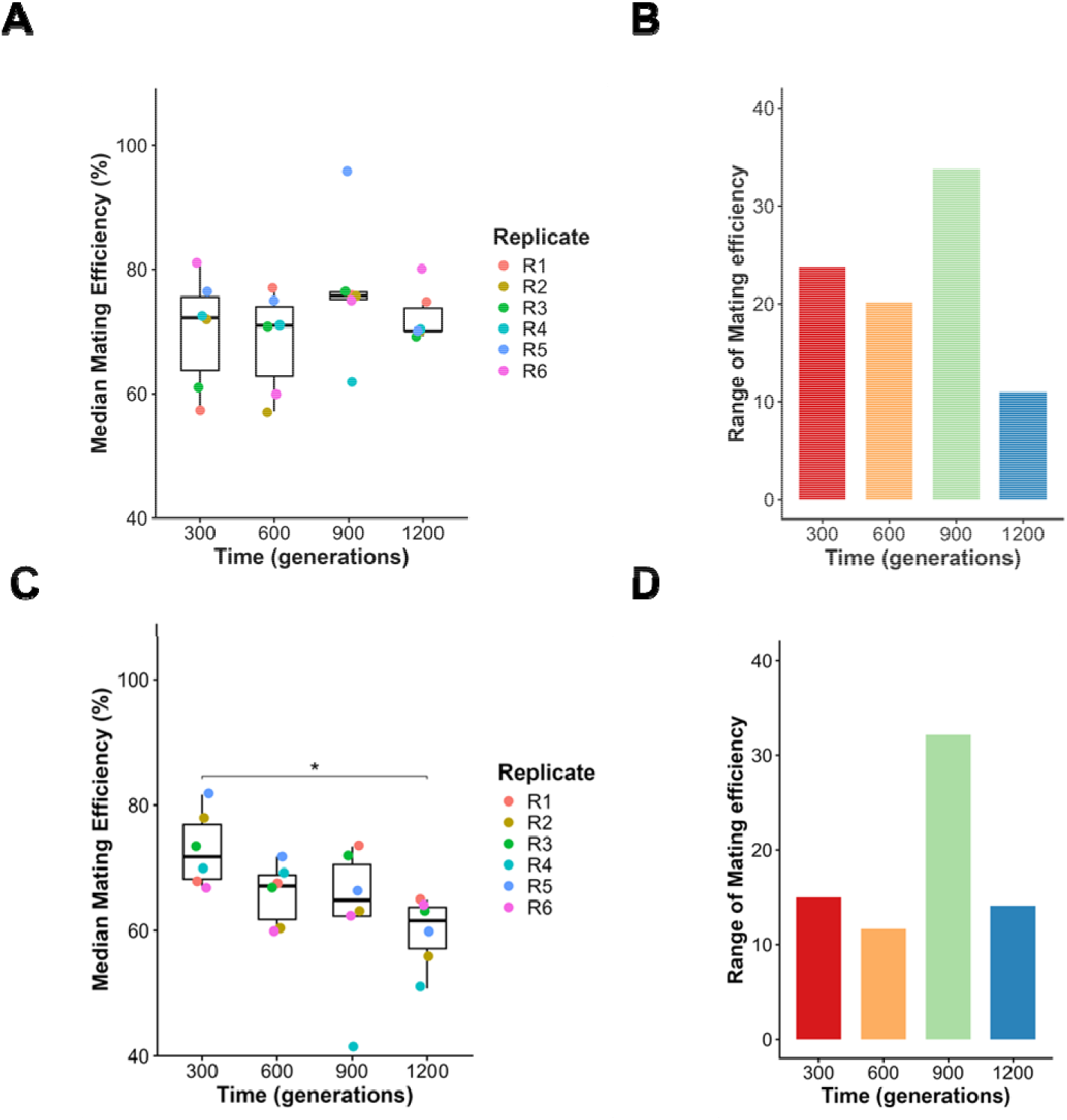
**(A)** Mating efficiency between glucose-evolved population and ancestor. The glucose-evolved population was sporulated, and haploids obtained from the evolved populations were crossed with ancestral haploids to assess intrinsic changes in mating ability. Median mating efficiency was calculated for every replicate at each time point. Median mating efficiency exhibited high variability among replicates at 300 and 600 generations, ranging from 55-85%. This variability decreased at later time points. However, no significant differences in mean median mating efficiency were observed across time (Kruskal Wallis test, p>0.05). **(B)** Range of the six median mating efficiency in glucose-ancestor hybrids is maximum at 900 generations and is the least at 1200 generations. **(C)** Mating efficiency between galactose-evolved population and ancestor. The galactose-evolved population was sporulated, and haploids obtained from the evolved populations were crossed with ancestral haploids to assess intrinsic changes in mating ability. Median mating efficiency was calculated for every replicate at each time point. Mating efficiency showed a gradual decline with time (Kruskal Wallis test, p<0.05), decreasing from 75-85% at 300 generations to 50-65% by 1200 generations (Dunn’s test, p<0.05). **(D)** Range of the six median mating efficiency in glucose-ancestor hybrids. It is maximum at 900 generations and is the least at 1200 generations.

In contrast, galactose-evolved populations exhibited a different pattern. Mating efficiency with the ancestor declined more gradually over time, from ∼80% at 300 generations to ∼60% by 1200 generations (**Figure 4C**). The decrease results in a statistically significant change over the 1200 generations, with the evolved lines mating with a significantly lower efficiency compared to the evolved lines earlier in the experiment. As seen in the Figure, the decrease in the mating efficiency is more or less linear with the number of generations. The six galactose-evolved lines exhibit approximately identical variance of the median mating efficiencies at the four time points (with one exception, i.e., line 4 at 900 generations) (**Figure 4D**). Note that the reduced ability of the galactose-evolved lines to mate with the ancestor is despite the evolved galactose-lines mating with near ancestor efficiency with themselves, or the glucose-evolved lines. Hence, the reduced mating efficiency is not a result of a compromised mating machinery in the cell.

These results indicates that evolved populations accumulate changes that affect their intrinsic ability to mate. The distinct temporal patterns observed in glucose- and galactose-evolved populations further suggest that the nature and dynamics of these intrinsic changes depend on the selective environment. These results are illustrated in **Supplement Figure S3**, where the within-line mating efficiency is often higher than the mating efficiency between the evolved lines and the ancestor.

### Genomic changes during adaptation

We previously genome sequenced all 12 lines at the four time points (300, 600, 900, and 1200 generations) and compared with the ancestor (48). In *S. cerevisiae*, the pathways regulating vegetative growth and mating are interconnected due to the shared MAPK signalling network (49, 50). Low nutrient availability suppresses activation of mating pathways as mating is a metabolically expensive process (51). During long-term evolution in nutrient-limited environments, mutations that alter mating could therefore be selectively favoured. In our work, however,, no such mutations were identified in the canonical mating genes or pheromone signalling pathways. Mutations were observed in genes such as *MSS11, MFG1, SNF2*. These genes are not direct components of mating pathway but are involved in regulation of filamentous and invasive growth in response to nutrient limitation.

*MSS11* is a transcriptional regulator that functions together with *FLO8* and *MFG1* to control expression of *FLO11*, required for filamentous or invasive growth, and also forms a part of conserved regulatory complex governing morphogenetic switching in yeast (52, 53). Similarly, *SNF2*, a component of SWI/SNF chromatin-remodeling complex, influences transcriptional programs involved in nutrient response (54). Although these genes are not directly involved in mating pathway, their disruption may indirectly influence mating behaviour due to alteration in growth. Since mating is a multi-step process involving pheromone production, pheromone sensing, adhesion, and cell fusion, small-effect mutations across multiple pathways may collectively alter mating behaviour. The reduction in mating efficiency can arise from cumulative regulatory changes rather than direct disruption of mating machinery.

Divergence of populations adapting to different environments may lead to emergence of pre-zygotic barriers. In the present study, there is no emergence of pre-zygotic barrier between glucose- and galactose-evolved populations. The glucose-evolved populations consistently exhibit higher fitness in glucose relative to the galactose-evolved populations and galactose-evolved populations exhibit a higher fitness in galactose when compared to the glucose-evolved populations, indicating progression towards environment specific adaptation. Although the differences are not statistically significant at every time point, the overall trend is consistent **(Supplement Figure S4A-B)**. Also, genomic analyses revealed that only a small fraction of SNPs were shared between glucose- and galactose-evolved populations, while majority of mutations – including fixed, segregating, and extinct variants – were unique to each environment **(Supplement Figure S4C)**. CNV analyses also showed that around 200 genes deleted in glucose-evolved populations were duplicated in galactose-evolved populations (48). Despite these genomic differences, no detectable pre-zygotic barriers emerge between glucose- and galactose-evolved populations across 1200 generations. Nonetheless, reduction in mating efficiency is observed when evolved populations are allowed to mate with the ancestor. This observation suggests that reproductive isolation may not depend on accumulation of genetic differences in mating pathway but depend on other traits that evolved in response to the environmental differences.

One possible explanation for reduced mating efficiency in evolved-ancestor crosses is due to the differences in mitotic growth rates of the evolved and ancestor haploids. Mating in *S. cerevisiae* progresses by arresting the cell cycle in G1 phase (55). Large differences in growth rate can create temporal mismatch and lead to lowered mating efficiency. Since, glucose- and galactose-evolved populations have comparable growth rates than any of the evolved population and ancestor (Supplement Figure S4A-B), there are significant reductions in mating efficiency in evolved-ancestor crosses than in glucose-galactose crosses. This implies that emergence of pre-zygotic barriers may not depend on alternation of mating pathways but on traits that allow temporal isolation. However, this interpretation is hypothetical and would require experimental validation.

## Discussion

Understanding how reproductive isolation emerges from evolving populations remains a central challenge in evolutionary biology (4, 5, 10). By tracking mating efficiency during prolonged asexual evolution, our study provides a window into the earliest stages of divergence in a facultative sexual organism (8, 12, 15, 16).

Our results show that mating traits evolve rapidly despite the absence of direct selection on mating. This indicates that mating efficiency is highly sensitive to mutations accumulated during adaptation, likely through pleiotropic effects on interconnected regulatory networks. The environment plays a critical role in shaping these outcomes: adaptation to galactose led to substantial perturbations in mating efficiency, whereas glucose-evolved populations remained comparatively stable. This suggests that the evolutionary fate of reproductive traits is tightly coupled to the ecological context and the specific targets of selection.

Early changes in mating compatibility are transient and non-monotonic, indicating that initial barriers to mating can arise rapidly but need not persist (8, 13, 17, 42). This contrasts with classical models that assume the steady accumulation of incompatibilities (56-61), and instead supports a view in which early divergence reflects reversible perturbations of underlying cellular networks, potentially stabilized or erased through compensatory processes (39, 40, 62, 63).

More broadly, our results suggest that early reductions in compatibility are not driven by a single mechanism, but emerge from the interplay between lineage-specific changes and interactions between diverging genomes. Critically, these effects appear insufficient, on their own, to produce stable reproductive isolation over short evolutionary timescales.

Finally, the pronounced divergence among replicate populations highlights the role of historical contingency in shaping reproductive traits (13, 19, 64). The emergence of incompatibilities is therefore not strictly predictable, but depends on the specific evolutionary paths taken by individual lineages, emphasizing the combined influence of stochastic processes and selection during the earliest stages of divergence (13, 14, 65, 66).

These findings have broader implications for our understanding of speciation. While long-term divergence may involve the accumulation of stable incompatibilities, our results suggest that the earliest stages are characterized by dynamic, reversible, and lineage-specific changes (3-5, 8, 14, 60). In facultative sexual organisms, where mating may be infrequent, reproductive traits may experience weak or indirect selection, allowing them to drift or respond pleiotropically to adaptation (30, 67-70). Under such conditions, transient barriers may arise without necessarily leading to long-term isolation (2, 8, 10, 12).

In summary, our study shows that asexual adaptation can rapidly generate changes in mating efficiency and compatibility, but these changes are heterogeneous, environment-dependent, and often reversible. Rather than a steady progression toward reproductive isolation, early divergence appears to involve fluctuating and contingent dynamics, shaped by both ecological context and lineage-specific evolutionary paths (71-74).

## Methods

### Adapative laboratory evolution and growth rate measurements

Twelve replicate populations of *S. cerevisiae* were evolved in 0.5% glucose and 0.5% galactose environments until 1200 generations. Growth rate measurements were performed in the respective evolution environments every 300 generations as described previously (48).

### Isolation of haploids for mating efficiency experiment

The evolved diploids were revived in their respective evolution environments, and sporulated to obtain haploids. And haploids from each environment were isolated based on the auxotrophic markers using the following protocol:

### Sporulation

5µL of saturated culture of evolved lines were used to make a patch on pre-sporulation media (6% glucose, 1% yeast extract, 2% peptone, 2% agar). It was incubated at 30ºC for 24 hours. Using the patch on pre-sporulation media, a fresh patch was made on the sporulation media (1% Potassium acetate, 0.12mg/mL complete amino acid mixture, 2% agar). The plate was incubated at 25ºC for 5 days (75).

### Ascospore isolation

After 5 days of incubation, ascospore isolation was performed using as described in (76) except that lyticase was used. Briefly, a portion of the patch of each line is taken from the sporulation media and resuspended in 500µL of autoclaved water in 1.5mL vials. 2.5 units of lyticase enzyme was added and incubated at 30ºC for 2 hours. 270mg of glass beads were added and samples were vortexed for 10 minutes. Dilutions were done and were plated on YPD. Plates were incubated for 36-48 hours at 30ºC until single colonies were obtained.

### Screening of haploids

The auxotrophic markers were used to screen for the haploids. The ‘a’ strain has *TRP1* gene and ‘⍰’ strain has *LEU2* gene inserted in the *ARS314* region near *MAT* locus on chromosome III. Hence, ‘a’ can grow on medium lacking tryptophan and ‘⍰’ can grow on medium lacking leucine and the diploid can grow on a medium that lacks both tryptophan and leucine. Single colonies obtained on the YPD plate were restreaked on trp^-^, leu^-^, trp^-^leu^-^ plates (2% glucose, 0.5mg/mL drop out amino acid mixture, 6.6mg/mL nitrogen base, 2% agar) to identify haploids.

### Mating efficiency experiment

Mating efficiency experiments were performed every 300 generations using the haploids obtained after sporulation. The mating efficiency of the following haploid combinations were determined:

- Within environment: glucose ‘a’ x glucose ‘⍰’ and galactose ‘a’ x galactose ‘⍰’
- Evolved hybrids: glucose ‘a’ x galactose ‘⍰’
- Evolved-ancestor hybrids: glucose ‘a’ x ancestral ‘⍰’, galactose ‘a’ x ancestor ‘⍰’
- Control: ancestor ‘a’ x ancestor ‘⍰’.

The haploids were inoculated in 5mL YPD and grown until saturation at 30ºC with 250rpm shaking. Following which, mating efficiency grids were setup on a YPD agar plate. The grid is a 1cm × 1.5cm rectangle divided into 3 boxes such that each has a dimension of 1cm × 0.5cm. 5µL of the YPD culture of the two mating types are laid on the extreme boxes. The plates were incubated for 24 hours at 30ºC. Then, equal number of cells of each haploid were mixed and laid down in the middle box and incubated at 30ºC for 7 hours. After the incubation, cells were scraped from the center box and diluted and plated on YPD to obtain isolated single colonies. The single colonies were then transferred onto the double drop out plate to identify the number of diploid colonies (36).

Mating efficiency was calculated as follows:

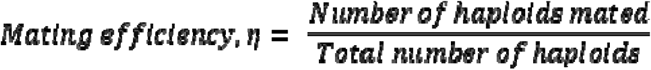

## Acknowledgements

This work was funded by DBT/Wellcome Trust (IA) via grant number IA/S/19/2/504632. AM was supported by the IRCC Internship Award, IIT Bombay. We thank all lab members for their feedback.

## Notes

### Competing Interest Statement

The authors have declared no competing interest.

