## Supplement Figures. for "Asexual Adaptation Drives Transient and Reversible Changes in Mating Efficiency"

**Running Title:** Asexual Adaptation Alters Mating Efficiency
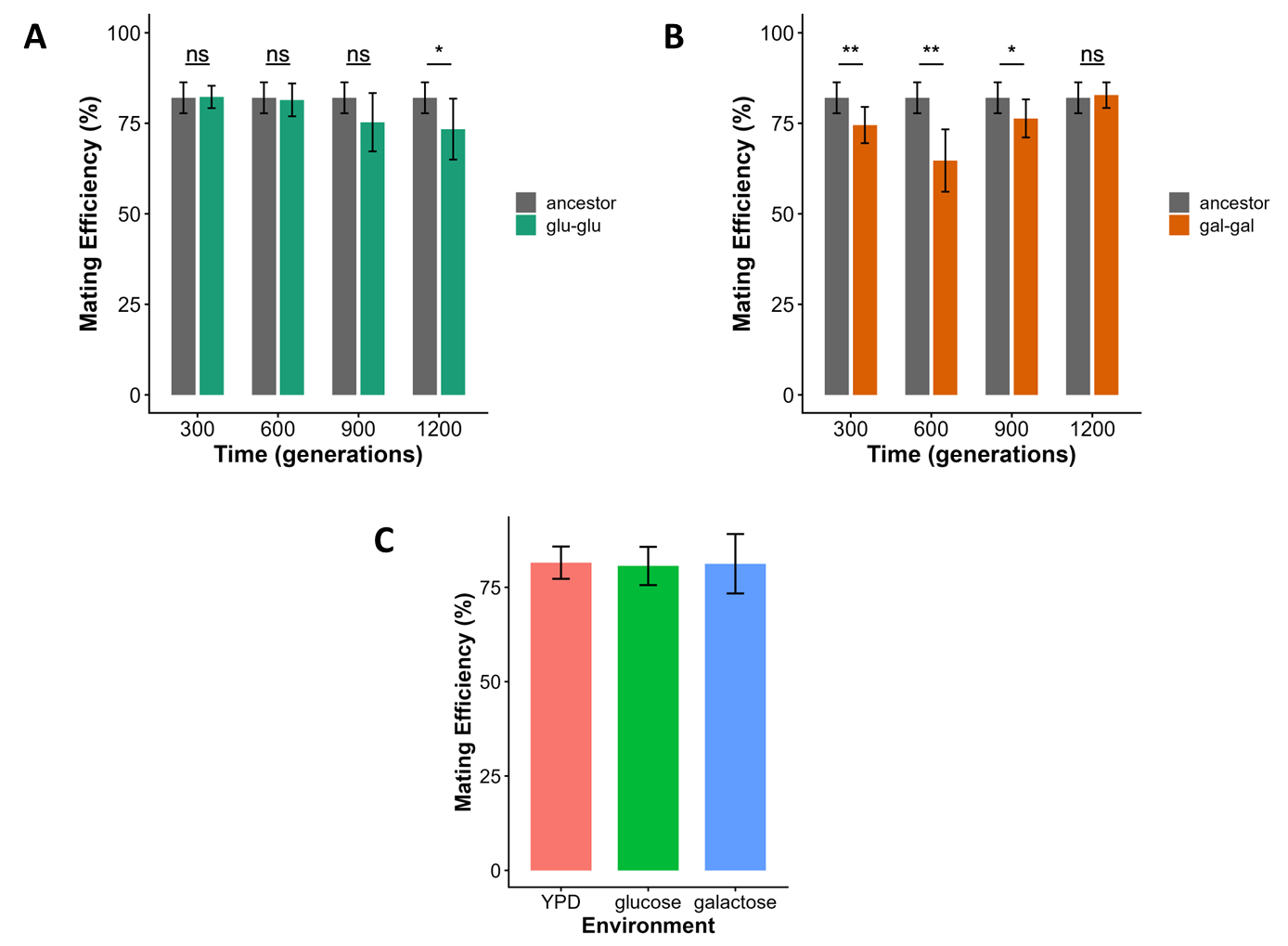

**Figure S1: Comparison of within-environment mating of glucose- and galactose-evolved populations with ancestral mating efficiency.** A significant difference in within-environment mating efficiency between glucose-evolved populations and the ancestor was observed only at 1200 generations **(A)**. In contrast, galactose-evolved populations showed significant differences at all time points except 1200 generations when compared with ancestor **(B).** Significance was determined using pairwise two-tailed Wilcoxon Mann Whitney test. The observed differences in galactose-evolved populations were not due to a physiological response caused by shifting galactose-evolved populations onto glucose-containing media for the mating assay, as the response is not consistent across time points, and preculturing ancestral populations in galactose did not show changes in mating efficiency. Mating efficiency of haploids remained constant irrespective of preculture environment **(C)**.

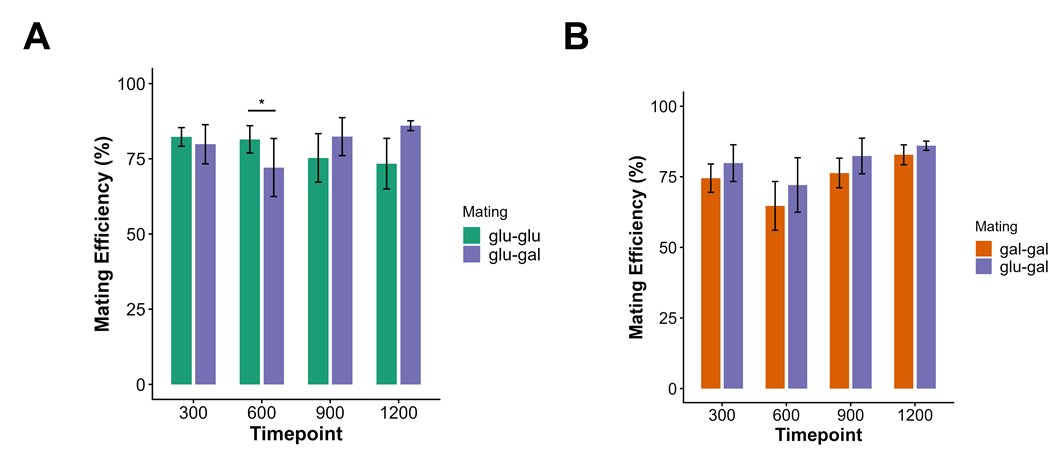

**Figure S2.** Comparison of glu-gal hybrids’ mating efficiency with respective within line mating efficiency at all four time points. The hybrid mating efficiency is either same or greater than its parent mating efficiency except at 600 generations when compared to glu-glu mating efficiency. (one-tailed Wilcoxon rank sum test, p<0.05 (*), p<0.01 (**), p<0.001(***)).

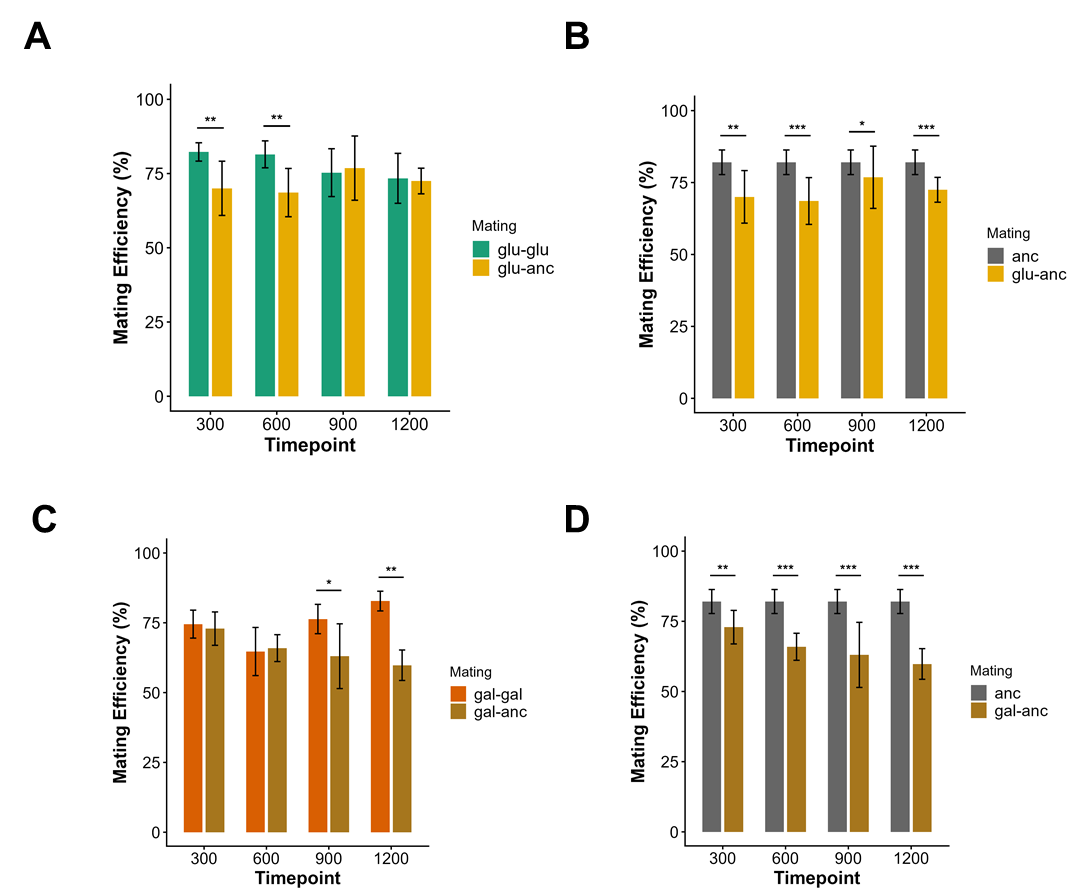

**Figure Supplement 3. (A-B)** Comparison of glucose-ancestor hybrids’ mating efficiency with respective parent populations at all four time points. The glu-anc hybrid mating efficiency is lower than glu-glu mating efficiency at early time points while it is lower than ancestral mating efficiency at all time points. Reproductive isolation occurs in the initial stage and is not maintained with time. **(C-D)** Comparison of galactose-ancestor hybrids’ mating efficiency with respective parent populations at all four time points. The gal-anc hybrid mating efficiency is lower than gal-gal mating efficiency at later time points while it is lower than ancestral mating efficiency at all time points. Reproductive isolation occurs at later stages of the experiment. (one-tailed Wilcoxon rank sum test, p<0.05 (*), p<0.01 (**), p<0.001(***))

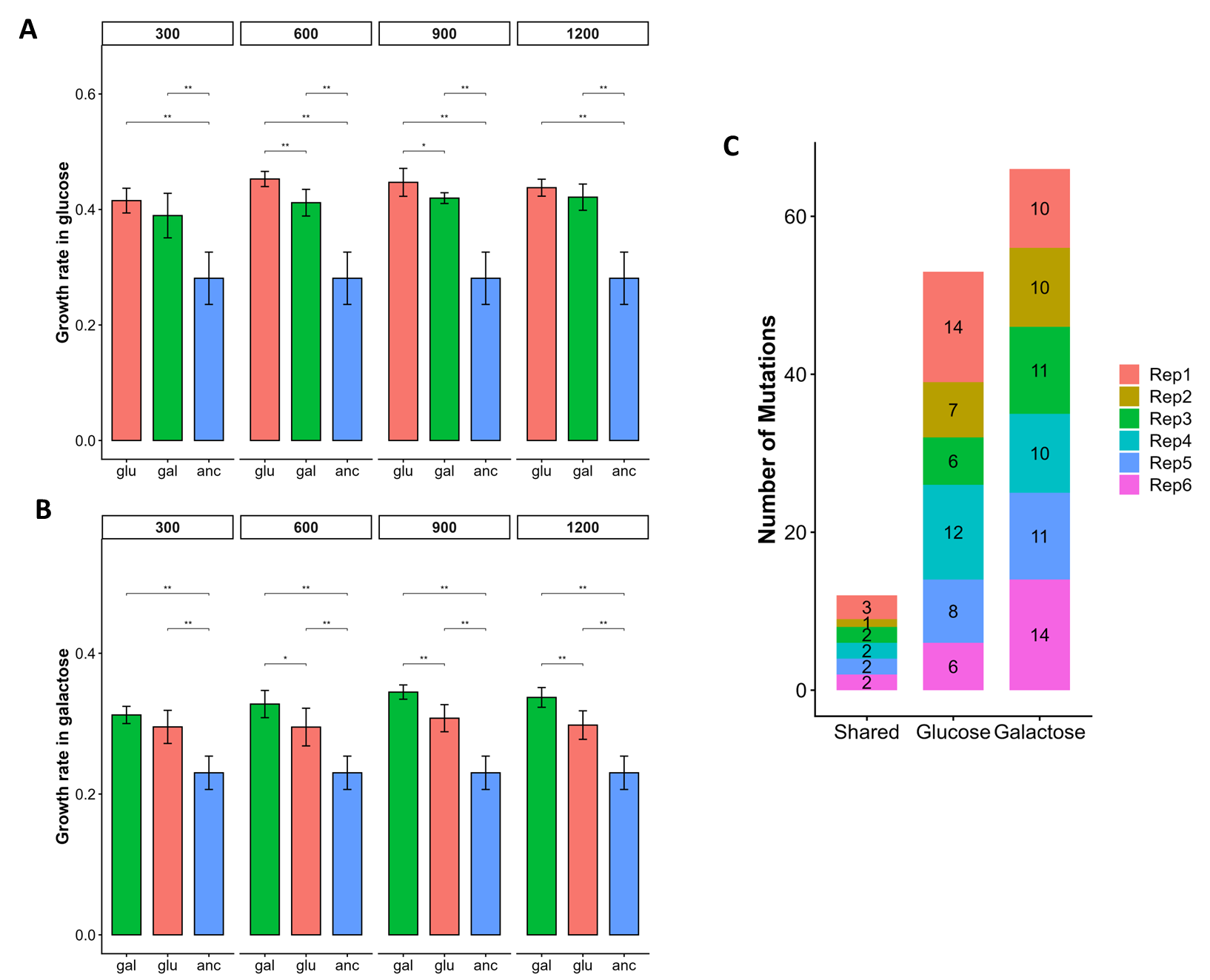

**Figure S4: Differences in glucose- and galactose-evolved populations.** Growth rate of glucose- and galactose-evolved populations were measured in both glucose and galactose. Glucose-evolved populations consistently have higher growth rate in glucose than galactose-evolved populations **(A)**. Similarly, galactose-evolved populations have higher growth rate in galactose than glucose-evolved populations **(B)**. Although these differences are not significantly different at every time point, the trend is consistent. Nonetheless, in both the environments and every time point, the evolved-populations’ growth rate is significantly higher than ancestral growth rate. **(C)** SNPs/Indels in the evolved populations were compared pairwise depending on the replicates that were allowed to mate in the glucose-galactose hybrid crosses. Glucose- and galactose-evolved populations have more unique mutations than shared mutations. Hence, the evolved populations have non-overlapping genomic targets.
